# Local nuclear to cytoplasmic ratio regulates H3.3 incorporation via cell cycle state during zygotic genome activation

**DOI:** 10.1101/2024.07.15.603602

**Authors:** Anusha D. Bhatt, Madeleine G. Brown, Aurora B. Wackford, Yuki Shindo, Amanda A. Amodeo

## Abstract

Early embryos often have unique chromatin states prior to zygotic genome activation (ZGA). In *Drosophila*, ZGA occurs after 13 reductive nuclear divisions during which the nuclear to cytoplasmic (N/C) ratio grows exponentially. Previous work found that histone H3 chromatin incorporation decreases while its variant H3.3 increases leading up to ZGA. In other cell types, H3.3 is associated with sites of active transcription and heterochromatin, suggesting a link between H3.3 and ZGA. Here, we test what factors regulate H3.3 incorporation at ZGA. We find that H3 nuclear availability falls more rapidly than H3.3 leading up to ZGA. We generate H3/H3.3 chimeric proteins at the endogenous H3.3A locus and observe that chaperone binding, but not gene structure, regulates H3.3 behavior. We identify the N/C ratio as a major determinant of H3.3 incorporation. To isolate how the N/C ratio regulates H3.3 incorporation we test the roles of genomic content, zygotic transcription, and cell cycle state. We determine that cell cycle regulation, but not H3 availability or transcription, controls H3.3 incorporation. Overall, we propose that local N/C ratios control histone variant usage via cell cycle state during ZGA.

## Introduction

Genome accessibility can be dynamically regulated through controlled incorporation of variant histones^1–3^. In most tissues, replication-coupled (RC) histones, produced during S-phase, generate the majority of nucleosomes^2,4,5^. RC histones have unusually high copy number, lack introns, and contain specialized UTRs to facilitate their rapid production during S-phase^3,6–9^. Conversely, replication-independent (RI), “variant” histones are made throughout the cell cycle and incorporated into specific genomic regions^4,10^. The exchange of RC and RI histones on chromatin is a common feature of early embryonic development especially during zygotic genome activation (ZGA)^11–17^. During ZGA, chromatin undergoes extensive remodeling to facilitate bulk transcription and establish heterochromatin^18–25^.

The pre-ZGA cell cycles in many organisms depend on maternally supplied components, including histones^26–29^. These cycles are unusual since they oscillate between S and M without growth phases, leading to an exponential increase in the nuclear to cytoplasmic (N/C) ratio^29–34^. The N/C ratio, in turn, controls the timing of cell cycle slowing and ZGA^31–33,35–37^. Titration of maternal histones against the increasing amount of DNA has been proposed to contribute to N/C ratio sensing in the early embryo^38–45^. Another hallmark of ZGA is histone variant exchange on chromatin. In many organisms, maternally supplied, embryonic-specific linker histone variants are replaced by RC H1s during ZGA^12–14,16,17^. Concurrently, the RC nucleosomal H2A is also replaced by RI H2Av as a consequence of the lengthened interphase in cycles leading up to ZGA in *Drosophila*^15,46^. Similarly, we have previously shown that RC H3 is replaced by RI H3.3 during these same cycles, though the cause remains unclear^29^.

H3.3 is essential for proper embryonic development in mice, *Xenopus*, and zebrafish^47–51^. In *Xenopus,* the H3.3-specific S31 residue is required for gastrulation while its chaperone binding site is dispensable^50^. In *Drosophila*, H3.3 nulls survive until adulthood using maternal H3.3 but are sterile^52^. Flies expressing H3 from the H3.3 enhancer generated conflicting results as to whether H3.3 protein or simply a source of replication-independent H3 is required for fertility^52,53^. The H3/H3.3 pair is particularly interesting during ZGA because H3.3 is enriched at sites of active transcription and in heterochromatin, which are both established during ZGA^2,4,54^.

Here, we examine the factors that contribute to H3.3 incorporation at ZGA in *Drosophila*. We identify a more rapid decrease in the nuclear availability of H3 than H3.3 over the final pre-ZGA cycles. We find that chaperone binding, not gene expression, controls incorporation patterns using H3/H3.3 chimeric proteins at the endogenous H3.3A locus. The increase in H3.3 incorporation depends on the N/C ratio. Since the N/C ratio affects many parameters of embryogenesis, we further test the contributions of genomic content, zygotic transcription, and cell cycle states. We identify cell cycle regulation, but not H3 availability or transcription, as a major determinant of H3.3 incorporation. Overall, we propose a model in which local N/C ratios regulate chromatin composition via cell cycle state during ZGA.

## Results

### The interphase nuclear availability of H3 decreases more rapidly than H3.3 over the pre-ZGA cycles

To understand in vivo dynamics of the H3/H3.3 pair during ZGA in *Drosophila*, we previously tagged H3 and H3.3 with a photo-convertible Dendra2 protein, (H3-Dendra2 and H3.3-Dendra2) at a pseudo-endogenous H3 locus and the endogenous H3.3A locus respectively (Figures S1A-B)^29^. *Drosophila* ZGA occurs after 13 rapid syncytial nuclear cycles (NCs) and is accompanied by cell cycle slowing and cellularization. We have previously shown that with each NC, the pool of free H3 is depleted and its levels on chromatin decrease (Figure S1C-D)^29^. In contrast, H3.3 chromatin levels increase during the same cycles (Figure S1C-D)^29^. To test if changes in the relative nuclear availability of H3 and H3.3 mirror the observed chromatin incorporation trends, we measured the nuclear intensities of H3-Dendra2 and H3.3-Dendra2 in each interphase. We observed that H3 nuclear intensities decreased by ~40% between NC10 and NC13 as previously shown (Figure 1A-B)^29^. However, when we measured H3.3-Dendra2 nuclear intensities we found that they decreased by only ~20% between NC10 and NC13 (Figure 1A-B). To further assess how nuclear uptake dynamics changed during these cycles, we tracked total nuclear H3 and H3.3 in each cycle and found that H3.3 accumulation reduced more slowly than H3 (Figure 1C-D).

**Figure 1:**
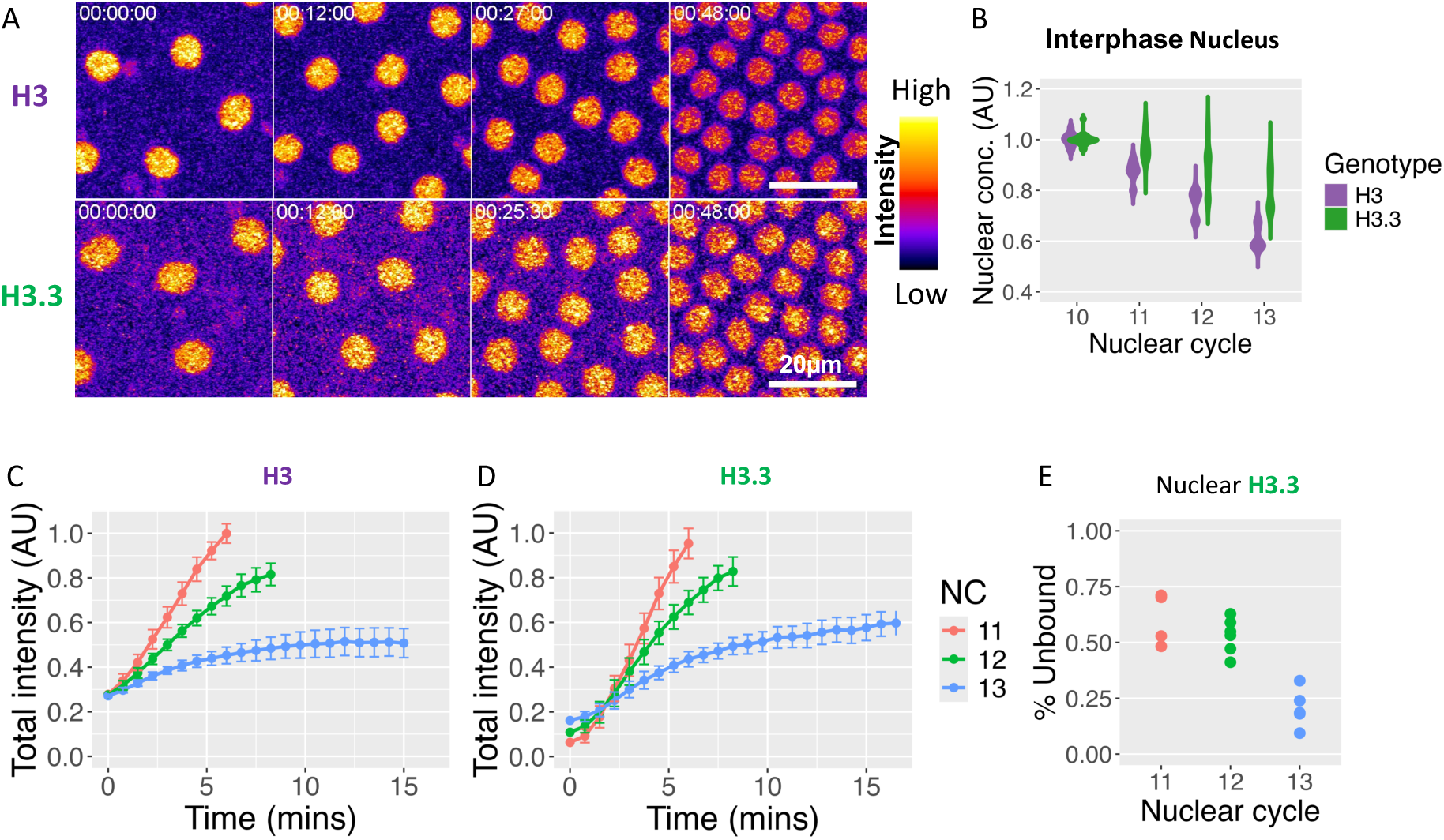
Interphase Nuclear availability of H3 decreases more rapidly than H3.3 over the pre-ZGA cycles. **(A)** Maximum intensity projections of H3-Dendra2 (top) and H3.3-Dendra2 (bottom) interphase nuclei 45 seconds before nuclear envelope breakdown (NEB) from NC10-13. Images are pseudo-colored with non-linear look-up tables where purple indicates low and yellow indicates high intensities. **(B)** Average interphase nuclear pixel intensities for H3-Dendra2 and H3.3-Dendra2 45 seconds before NEB in NC10-13, normalized to the average individual NC10 values. H3 and H3.3 concentrations decrease over time, but H3 loss is relatively more rapid. **(C-D)** Summed (total) pixel intensities for each nucleus over time for NC11-13 normalized to the maximum NC11 values for H3-Dendra2 **(C)** and H3.3-Dendra2 **(D)**. Nuclear import plateaus after the first 5 mins for H3, but merely slows and does not plateau for H3.3 in NC13. **(E)** The fraction of photoconverted unbound H3.3-Dendra2 after NEB in NC11-13 (see materials and methods for details). The “free” pool of H3.3 falls with each cycle. (n=3 H3 and 5 H3.3 embryos in B-D and >= 5 embryos in F; Statistical comparisons for B can be found in Supplemental table 2).

The reduction in nuclear accumulation could be due to a decrease in nuclear import, an increase in nuclear export, or both. To test these possibilities, we quantified the rate of nuclear export by photo-converting Dendra2 during interphase and measuring red Dendra2 signal over time. Using this method, we have previously shown that nuclear export of H3 is negligible^29^. Here, we find that export of H3.3 is also negligible (Figure S1E). These data suggest that the distinct dynamics of H3 and H3.3 nuclear availability are due to their import dynamics. Though the change in initial import rates between NC10 and NC13 are similar between the two histones (Figure S1F), we observed a notable difference in their behavior in NC13. H3 nuclear accumulation plateaus ~5 minutes into NC13, whereas H3.3 nuclear accumulation merely slows (Figure 1C-D). These changes in nuclear import and incorporation result in a less dramatic loss of the free nuclear H3.3 pool than previously seen for H3 (Figure 1E)^29^.

### Chaperone binding sites regulate the differences in H3 and H3.3 chromatin incorporation

We next investigated what differences between H3 and H3.3 caused the observed trends in chromatin incorporation. There are two major differences between H3 and H3.3: protein sequence and expression pattern. H3 differs from H3.3 by four amino acids which create an additional phosphosite in H3.3 and generate differing affinities for specific H3-family histone chaperones^55^. H3 is also generally expressed at much higher levels and in a replication-dependent manner. To determine which factor controls nuclear availability and chromatin incorporation, we genetically engineered flies to express Dendra2-tagged H3/H3.3 chimeras at the endogenous H3.3A locus. These chimeras include (i) H3.3’s phosphosite replaced with Alanine from H3 (H3.3^S31A^) (ii) H3.3’s chaperone binding domain replaced with H3’s (H3.3^SVM^), and (iii) all four H3.3-specific amino acids replaced with those of H3 (H3.3^ASVM^), (Figure 2A). In all cases, the gene structure, including the promoter, intron, and UTRs of H3.3, remained intact and no other codons were changed to maximize similarity to the endogenous H3.3A locus.

**Figure 2:**
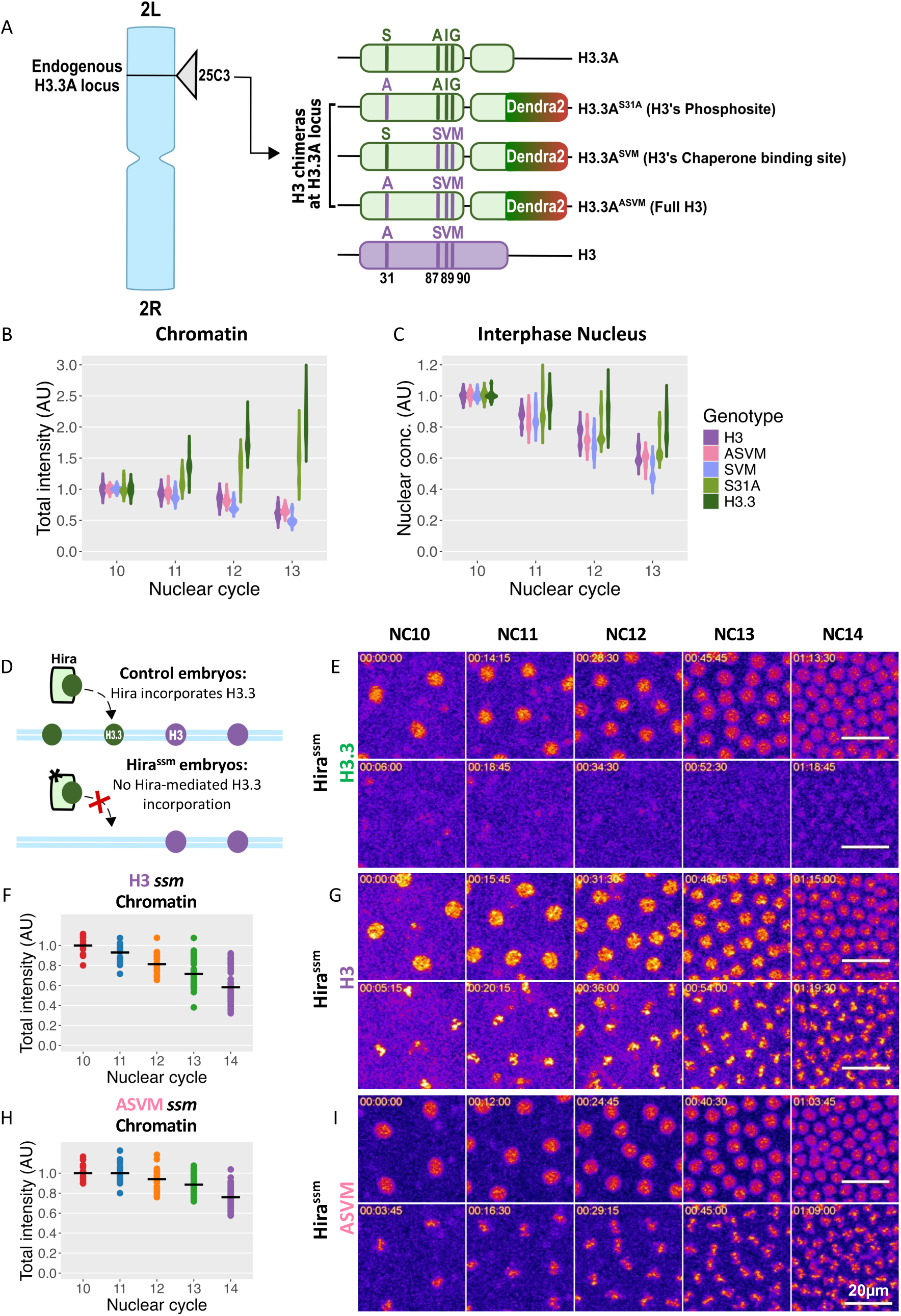
The chaperone binding site determines H3 variant chromatin incorporation. **(A)** Schematic of the Dendra2 tagged H3/H3.3 replacement chimeras at the endogenous H3.3A locus. S31A: H3.3 phosphosite (S) replaced with that of H3 (A), SVM: H3.3 chaperone binding site (AIG) replaced with that of H3 (SVM), and ASVM: all H3.3-specific amino acids replaced with those from H3. **(B)** Total intensities on mitotic chromatin of chimeras during NC10-13 normalized to their NC10 values. The same data for H3-Dendra2 and H3.3-Dendra2 are shown in Figure S1C. H3.3^S31A^ increases similarly to H3.3 while the constructs containing the H3 chaperone binding site decrease similarly to H3. **(C)** Interphase nuclear concentrations of chimeras 45 seconds before NEB during NC10-13 normalized to their NC10 values. H3-Dendra2 and H3.3-Dendra2 from Figure 1B included for reference. As seen for chromatin, nuclear accumulation generally follows the behavior of the chaperone binding site. **(D)** Schematic of H3.3 incorporation in control embryos and Hira^ssm^ mutants. H3.3 is imported to the nucleus, but the mutant Hira chaperone fails to incorporate H3.3. Hira mutants develop as haploids and undergo one additional fast nuclear division. **(E, G, I)** Representative maximum intensity projections during interphase and mitosis over NC10-14: interphase nuclei (top) and mitotic chromatin (bottom) for H3.3-Dendra2 **(E)**, H3-Dendra2 **(G)**, and H3.3^ASVM^-Dendra2 **(I)**. Images are pseudo-colored with non-linear look-up tables such that purple indicates low intensities and yellow indicates high intensities. H3 and H3.3^ASVM^ continue to accumulate on chromatin in the absence of Hira. **(F, H)** Total intensities of H3-Dendra2 **(F)** and H3.3^ASVM^-Dendra2 **(H)** on mitotic chromatin in Hira^ssm^ embryos between NC10-14 normalized to their average NC10 values. Though H3.3^ASVM^ is successfully incorporated without active Hira, the chromatin amounts decrease more slowly than H3. (n=5 all chimeras, 3 H3 ssm, 4 H3.3 ssm, and 5 H3.3^ASVM^ ssm embryos. Statistical comparisons for B and C can be found in Supplemental tables 6-7).

To study how chromatin incorporation differed in these chimeras, we measured their total intensities on mitotic chromatin during each nuclear cycle. We observed that, though H3.3^S31A^ chromatin incorporation was significantly reduced compared to H3.3 by NC13, its levels increased on chromatin over the nuclear cycles, resembling H3.3 more than H3 (Figure 2B, S2A). Conversely, the total amount of H3.3^SVM^ and H3.3^ASVM^ on mitotic chromatin fell over the nuclear cycles, similar to H3 (Figure 2B, S2B-C). This suggests that chromatin incorporation is mainly determined by the chaperone binding site. These results are broadly consistent with the final interphase nuclear concentrations and import dynamics where H3.3^S31A^ was intermediate between H3 and H3.3 while H3.3^SVM^ and H3.3^ASVM^ were more similar to H3 (Figure 2C, S2A-G). However, both nuclear H3.3^S31A^ and H3.3^SVM^ fell more quickly than H3.3 and H3 respectively, suggesting that chimeric histones may not be as stable and/or efficiently imported as their canonical counterparts. Together, these data indicate that the specific amino acid sequence of the chaperone binding site is the primary factor in differentiating the two histones for chromatin incorporation and nuclear import dynamics.

### H3 chaperone binding site conveys independence from Hira for chromatin incorporation

Since the chromatin incorporation of the H3/H3.3 chimeras appears to depend on their chaperone binding sites, we asked whether they still required the canonical H3.3 chaperone, Hira. We used Hira^ssm-185b^ (hereafter Hira^ssm^) flies which have a point mutation in the Hira locus^56^. This mutant Hira protein can bind but not incorporate H3.3 into chromatin (Figure 2D-E, S2H), resulting in sperm chromatin decondensation defects. These embryos develop as haploids and undergo one additional syncytial division before ZGA (NC14)^56^. The fall in nuclear concentration of H3 is slightly more gradual in the haploid Hira^ssm^ embryos than in wildtype, though H3 chromatin incorporation is not disrupted (Figure 1B, 2F-G, S2I). To test if H3-like chimeras expressed from the H3.3A locus use the canonical Hira pathway, we measured import and chromatin incorporation of H3.3^ASVM^ in Hira^ssm^. We found that H3.3^ASVM^ interphase nuclear concentration was more stable than H3 or H3.3 in Hira^ssm^ embryos (Figure 2G, S2J). This stability is reflected in H3.3^ASVM^ chromatin incorporation where it only drops by ~20% between NC10 and NC14 compared to the observed ~40% drop in H3 (Figure 2H-I). These data indicate that H3.3^ASVM^ chromatin incorporation is Hira independent, even when expressed from the H3.3A locus.

### Local N/C ratios determine H3 and H3.3 chromatin incorporation

Since the N/C ratio controls many aspects of pre-ZGA development we asked whether the local N/C ratio determines histone chromatin incorporation within a nuclear cycle. To test this, we employed mutants in the gene Shackleton (Shkl) whose embryos have non-uniform nuclear densities across the anterior/posterior axis (Figure 3A-B, Movie 1-2)^57^. In these embryos, impaired cortical migration of early nuclei increases the N/C ratio in the center and decreases it in the posterior, which results in frequent partial extra divisions at the posterior pole (Figure 3B, G)^57^. For our analyses, we manually defined low and high nuclear density regions, with the low-density region always undergoing an extra division (Figure 3B, see methods). To control for potential positional effects, we measured chromatin incorporation at the middle and pole regions of control embryos for comparison (Figure 3A). In control embryos, the drop in the total amount of H3 and rise in total H3.3 on chromatin are comparable between the middle and pole over the pre-ZGA cycles (Figure 3C-D). In contrast, in Shkl embryos, we observe decreased incorporation of H3 on chromatin at high nuclear densities compared to low nuclear densities (Figure 3E, S3A). This trend is reversed for H3.3, where chromatin from high density regions has more total H3.3 than chromatin from low density regions (Figure 3F, S3B).

**Figure 3:**
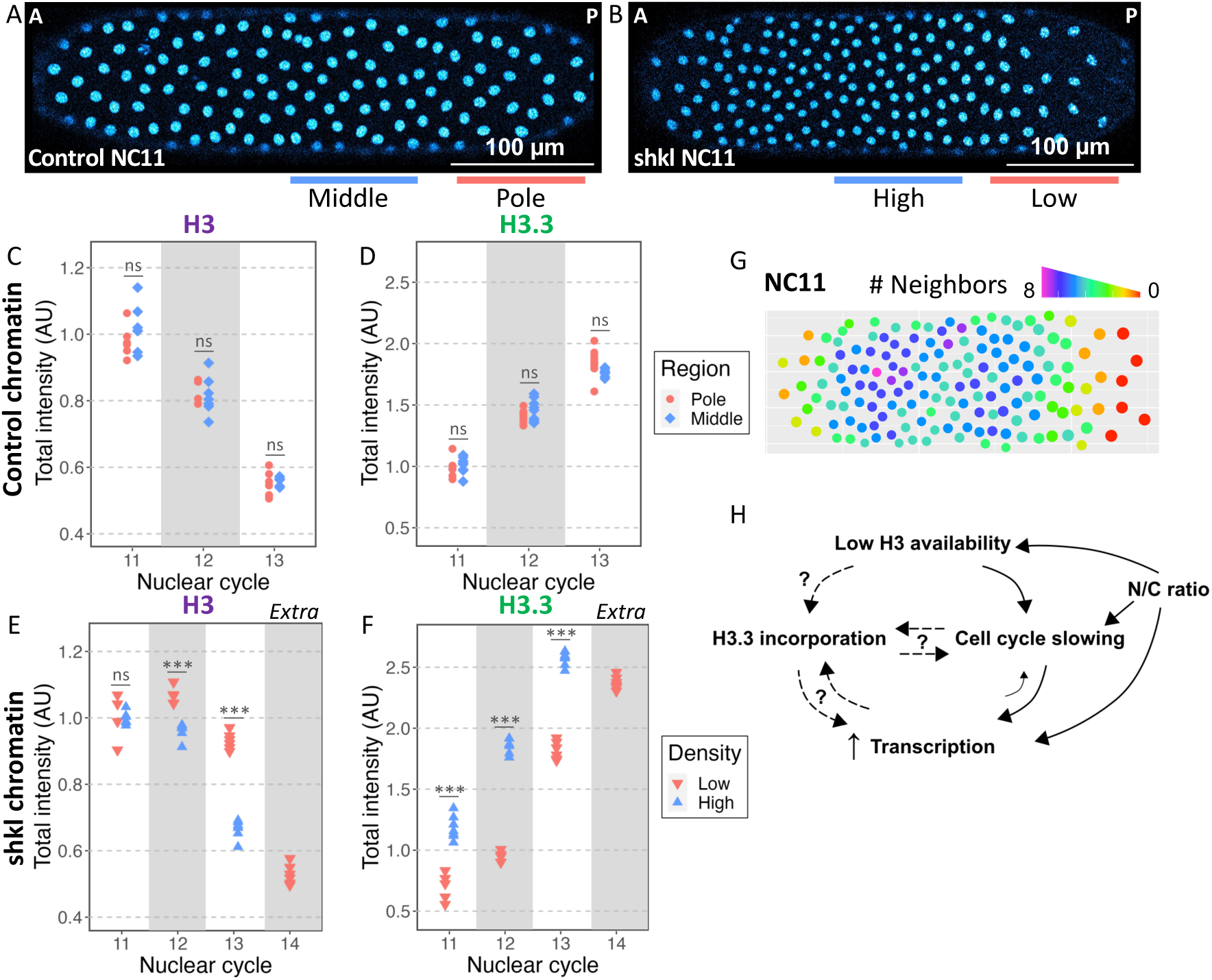
Local N/C ratios determine H3 and H3.3 chromatin incorporation. **(A)** Example NC11 control embryo with middle (blue) and pole (red) regions labeled as used in C and D. **(B)** Example NC11 shkl embryo with high (blue) and low (red) density regions labeled as used in E and F. **(C-D)** Total intensities on mitotic chromatin of H3-Dendra2 **(C)** and H3.3-Dendra2 **(D)** during NC11-13 in a representative control embryo where each point indicates a single nucleus. Total H3-Dendra2 intensities fall and H3.3-Dendra2 intensities rise uniformly between middle and pole regions within each cycle. **(E-F)** Total intensities on mitotic chromatin of H3-Dendra2 **(E)** and H3.3-Dendra2 **(F)** during NC11-13 in a representative shkl embryo where each point indicates a single nucleus. NC14 represents a partial extra division in the low-density region. Chromatin in the low-density region retains more H3 and incorporates less H3.3 within the same cell cycle compared to the high-density region. Similar results were observed in replicate embryos (Figure S3A-B). **(G)** Gradient in the number of neighbors for each nucleus at its minimum volume within a 20µm radius for the shkl embryo shown in B. **(H)** Direct and indirect mechanisms of H3.3 incorporation in response to the N/C ratio. H3.3 incorporation could be a direct result of reduced nuclear H3 availability. Here, the increasing demand for nucleosomes with the increasing numbers of genomes would be met by H3.3. The N/C ratio also controls transcription and cell cycle duration. H3.3 incorporation could be downstream of either process. (Statistical significance was determined by 2-way ANOVA, ns= p>.05, *** = p<0.001).

This observation indicates that incorporation of H3 and H3.3 are reciprocal and depend on the local N/C ratio leading to several possible models (Figure 3H). First, the H3 pool available for chromatin incorporation may become limiting at high N/C ratios leading to increased H3.3 incorporation. Second, since H3.3 is known to be associated with sites of active transcription^50,52,58–62^, the increased H3.3 incorporation might be downstream of N/C ratio dependent ZGA. Finally, since H3 is usually incorporated only during S-phase the changing H3 to H3.3 incorporation rates may be the result of N/C ratio-dependent cell cycle changes. Note, that all these processes feedback onto one another such as cell cycle slowing allowing time for ZGA^37,63^.

### H3 nuclear availability depends on the local N/C ratio

To ask whether nuclear availability can explain the N/C ratio-dependent differences in H3 and H3.3 incorporation, we measured their interphase accumulation in Shkl embryos (Figure 4A). Since H3 and H3.3 both have negligible nuclear export, their nuclear availabilities are determined by their import rates (Figure 1B, S1E)^29^. To assess the impact of the N/C ratio on nuclear import in individual nuclei, we calculated the number of neighbors within a 20 µm radius for each nucleus at its minimum volume (Figure 3G, Figure S3C). We then binned the nuclei by their number of neighbors and determined their nuclear import curves for both H3 and H3.3. In control NC13 embryos, there is little variation in the number of neighbors and all nuclei import H3 and H3.3 similarly (Figure 4B-E). In NC13 Shkl embryos, H3 import is anticorrelated with the local N/C ratio (Figure 4F, H, S3D). We observed slower H3 nuclear uptake at high N/C ratios resulting in lower total interphase H3 accumulation (Figure 4F). This was also reflected in the initial H3 import rates where the nuclei with fewer neighbors had higher slopes (Figure 4H). H3.3 uptake was less affected by the local N/C ratio (Figure 4G, I, S3E). A similar trend was also observed in NC12 for both histones, where more neighbors correspond to slower import. However, the range of behaviors was not as large as seen in NC13 (Figure S3F-G). These observations support a model where H3 pools are exhausted by the increasing N/C ratio, increasing the relative availability of H3.3 to H3 over the pre-ZGA cycles.

**Figure 4:**
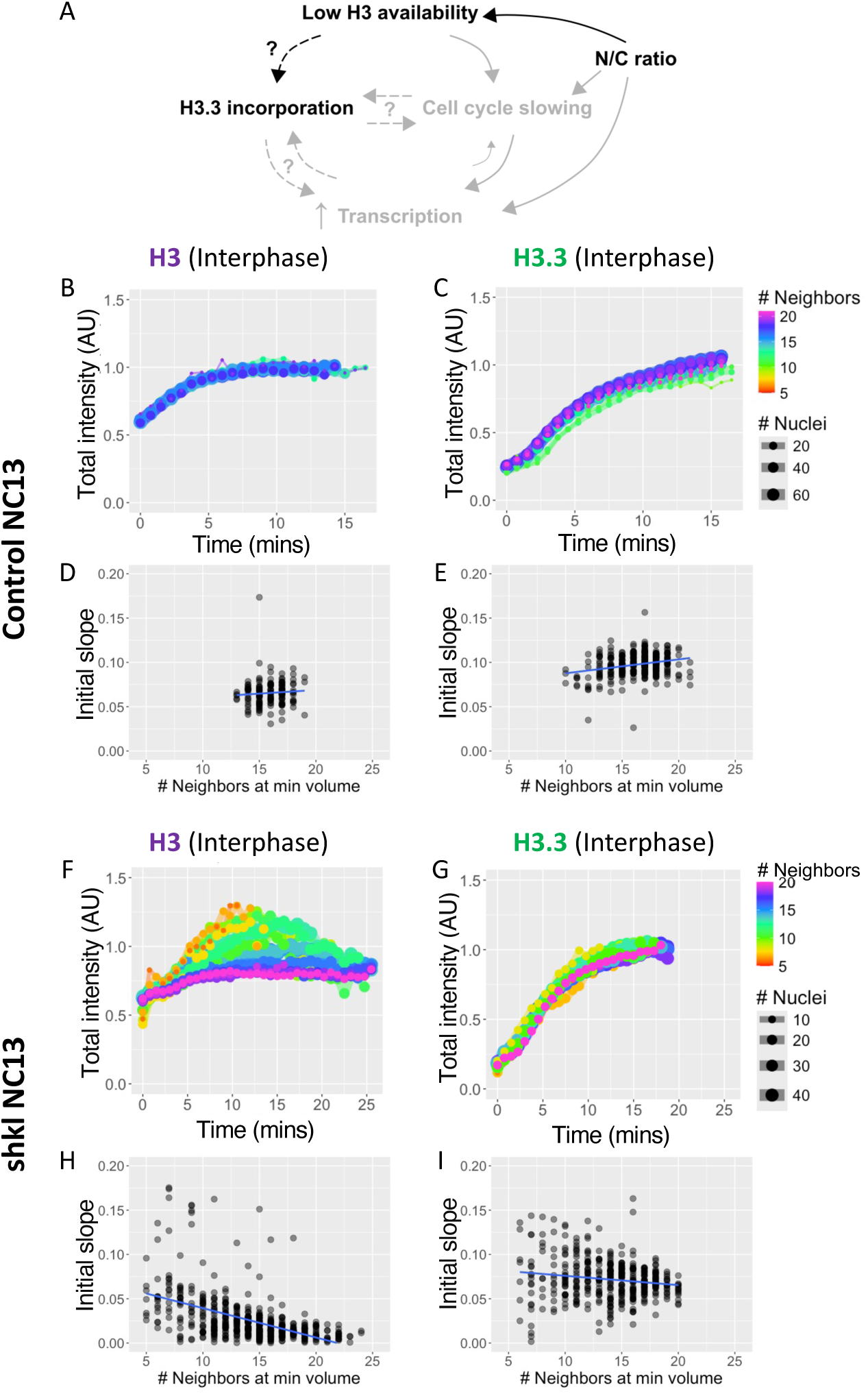
Local N/C ratios differentially affect H3 and H3.3 nuclear availabilities. **(A)** Schematic of how the N/C ratio might affect H3 and H3.3 chromatin incorporation through loss of available H3. Bolded portion is the hypothesis under consideration. **(B-C)** Total intensities over time for nuclei in representative NC13 control embryos binned by the number of neighbors as in 3G for H3-Dendra2 **(B)** and H3.3-Dendra2 **(C)**. Total intensity was normalized to the average maximum intensities achieved in NC13 and line color represents the number of neighbors. In controls there is little variation in the number of neighbors or the import of H3 and H3.3 across the length of the embryo. **(D-E)** Initial slopes of nuclear import curves from representative NC13 control embryos from B and C for H3-Dendra2 **(D)** and H3.3-Dendra2 **(E)** plotted by the number of nuclear neighbors. Note the uniformity in the number of neighbors and similarity in nuclear import behaviors in control embryos. **(F-G)** Total intensities over time for nuclei in representative NC13 shkl embryos binned by the number of neighbors as in 3G for H3-Dendra2 **(F)** and H3.3-Dendra2 **(G)**. Nuclear import and accumulation of H3 inversely correlate with the number of neighbors, suggesting H3 nuclear import is N/C ratio sensitive. H3.3 nuclear import is less N/C ratio sensitive than H3. Similar results were observed in replicate embryos (Figure S3D-G). **(H-I)** Initial slopes of nuclear import curves from representative NC13 shkl embryos from F and G for H3-Dendra2 **(H)** and H3.3-Dendra2 **(I)** plotted by the number of nuclear neighbors. The slopes reflect a faster H3 uptake in nuclei with fewer neighbors and a slower H3 uptake in nuclei with more neighbors. Slopes in some nuclei with more neighbors are near zero indicating that very little additional H3 is imported after nuclear envelope formation. Though the slopes reduce with the number of neighbors for H3.3, there is a non-negligible H3.3 import in the nuclei with the largest number of neighbors.

### H3.3 incorporation is not caused by exhaustion of H3 pools

Given that the available H3 seems to be depleted by the increasing N/C ratio we sought to test if H3.3 chromatin incorporation depends on the size of the H3 pool (Figure 5A). We hypothesized that as the embryo exhausted the supply of RC H3 it might increase the use of RI H3.3 to compensate. We knocked down Stem-loop binding protein (Slbp), which specifically binds and stabilizes the mRNAs of RC histones, including H3, but does not interact with H3.3 mRNAs^7,64,65^. Slbp RNAi dramatically decreases the size of the available H3 pool and results in frequent chromosomal segregation defects (Figure S4A)^39^. For this reason, we only analyzed embryos that appeared reasonably healthy until the final cell cycle under consideration. In embryos that survives through at least NC12, we found that H3.3 incorporation is largely unaffected by the reduction in RC H3 (Figure 5B). To further validate that the lack of effect on H3.3 incorporation was not due to inefficient Slbp-knockdown, we also tested H3.3 incorporation in embryos that already display severe bridging in NC11. In these embryos, we detected no difference in the H3.3 incorporation in NC10 mitosis (Figure S4B). These results strongly indicate that simply running out of H3 is not the cause of the observed increase in H3.3 on chromatin.

**Figure 5:**
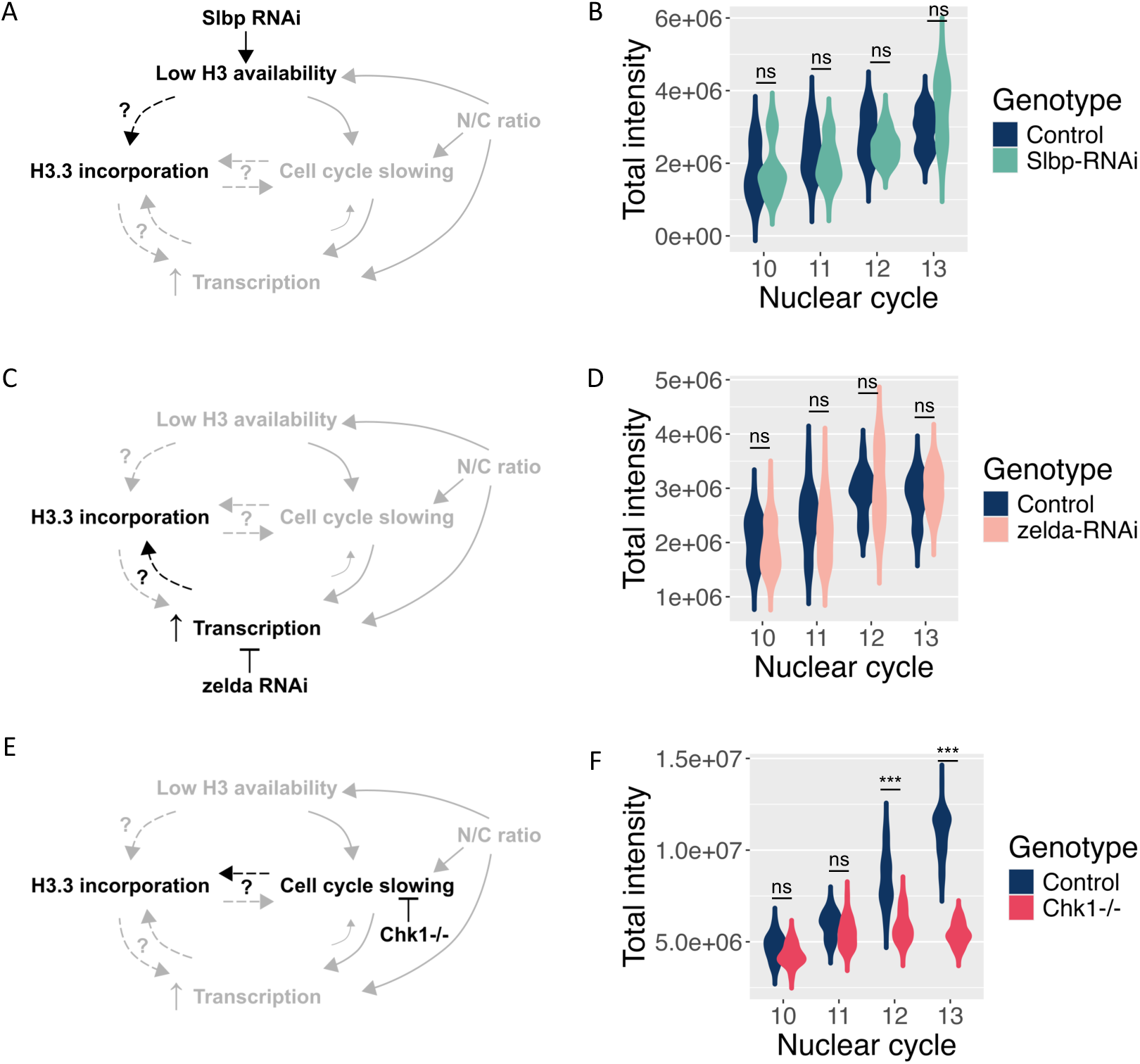
H3.3 incorporation depends on cell cycle state but not H3 availability or zelda-dependent transcription. **(A,C,E)** Schematics of different parameters that may regulate H3.3 chromatin incorporation. Bolded portion is the hypothesis under consideration in B, D and F respectively. **(A)** Slbp-RNAi decreases the size of the available H3 pool. **(B)** Total intensities of H3.3-Dendra2 on mitotic chromatin in white-RNAi (control) and Slbp-RNAi backgrounds during NC10-13. H3.3 incorporation does not increase upon lowering H3 availability. Note that most Slbp-RNAi embryos are arrested in NC13 without dividing and therefore do not contribute to the mitotic NC13 data. **(C)** zelda-RNAi inhibits the majority of zygotic transcription allowing us to test if H3.3 incorporation depends on transcription. **(D)** Total intensities of H3.3-Dendra2 on mitotic chromatin in white-RNAi (control) and zelda-RNAi backgrounds during NC10-13 normalized to their NC10 values. H3.3 incorporation does not change upon inhibiting zelda-dependent transcription. **(E)** Chk1 (grp^1^) mutation prevents cell cycle slowing allowing us to test if H3.3 incorporation is dependent on cell cycle state. **(F)** Total intensities of H3.3-Dendra2 on mitotic chromatin in control and embryos from chk1^−/−^ mothers during NC10-13. H3.3 incorporation is reduced in both NC12 and NC13 indicating that cell cycle state though not cell cycle duration regulated H3.3 incorporation. Note that these embryos are homozygous for H3.3-Dendra2 and have double the fluorescent intensity compared to all previous embryos. (n≥5 embryos, Statistical significance was determined by 2-way ANOVA, ns= p>.05, *=p<.05, **=p<0.01, *** = p<0.001)

### H3.3 incorporation does not depend on zelda-dependent ZGA

Since H3.3 is associated with sites of active transcription in other systems^50,52,58–62^, we next sought to test if H3.3 incorporation during ZGA depends on transcription. To do this, we knocked down the critical pioneer transcription factor zelda (Figure 5C). zelda controls the transcription of the majority of Pol II genes during ZGA and when zelda is disrupted, Pol II relocates to the histone locus body^66–68^. We found that H3.3 chromatin incorporation did not change in zelda RNAi embryos despite their inability to cellularize and longer NC13s (Figure 5D and S4C). This suggests that the large increase in H3.3 incorporation that we detect by microscopy in the final nuclear cycles does not depend on bulk ZGA.

### H3.3 incorporation depends on cell cycle state, but not cell cycle duration

Finally, to test the contribution of the cell cycle on the N/C ratio dependent accumulation of H3.3 on chromatin we used mutants in Chk1 (grapes in *Drosophila*) that are less efficient in cell cycle slowing (Figure 5E, S4C). These mutants have an unusually rapid NC13 and attempt to enter mitosis before their DNA is fully replicated resulting in mitotic catastrophe^69^. We found that H3.3 accumulation is disrupted as early as NC12 (P-value=10^−8^) in Chk1 mutants (Figure 5F). Importantly, the Chk1 mutants have relatively normal NC12 durations^69,70^. In our experiments, Chk1 NC12 was only ~1 minute faster than wildtype and Chk1 embryos with comparable cell cycle durations still displayed reduced H3.3 incorporation (Figure 5F and S4C). To further isolate the effect of cell cycle length on H3.3 incorporation we used the natural variation in NC13 duration in control embryos. When we plotted H3.3 chromatin signal against the total NC13 duration for control embryos we found no correlation (Figure S4D). This result suggests that cell cycle duration as such does not directly regulate H3.3 chromatin incorporation. Instead, Chk1 appears to regulate H3.3 incorporation in a manner that is not mediated solely by lengthening the cell cycle.

## Discussion

We demonstrate that H3.3 replaces H3 on chromatin leading up to ZGA in *Drosophila*. This process depends on the specific H3.3 chaperone binding site and is controlled by the N/C ratio. We tested which aspects of the N/C ratio control the dynamic incorporation of H3.3 and found that cell cycle state, but not H3 availability or bulk transcription, is the major regulator of H3.3 behavior. Chk1 mutants decrease H3.3 incorporation even before the cell cycle is significantly slowed. Cell cycle slowing has been previously reported to regulate the incorporation of other histone variants in *Drosophila*^15^. However, our results indicate that cell cycle state and not duration per se, regulates H3.3 incorporation. We speculate that this may be due to changes in chromatin state as a result of Chk1 activity. Late replicating regions and heterochromatin first emerge during ZGA, and Chk1 can control origin firing in many contexts^21,23,24,71,72^. Since H3.3 is often associated with heterochromatin, the decreased H3.3 incorporation in Chk1 mutants may be an indirect result of increased origin firing and decreased heterochromatin formation^71,72^. Another possibility is that the additional Chk1 phosphosite that is found in H3.3-S31 may be important for promoting H3.3 incorporation during ZGA^50^.

The interaction between H3-type histones and Chk1 has additional significance since H3 nuclear concentration has been proposed to directly regulate cell cycle length through H3 interactions with Chk1^40^. In Hira^ssm^ embryos that undergo one extra division before cell cycle slowing, the fall in nuclear H3 concentration between NC10 and the final fast cell cycle is strikingly similar to that seen in wildtype. Moreover, H3 nuclear concentrations appear to be strongly sensitive to the local N/C ratio in Shkl embryos. Together these data are consistent with a model in which H3 nuclear concentrations regulate cell cycle slowing. However, H3.3 nuclear concentrations are less sensitive to the local N/C ratio than H3. Since H3.3 has an additional Chk1 phosphorylation site compared to H3 it may have different regulatory interactions with Chk1^50,73^. The relative contributions of both H3 and H3.3 nuclear availability to cell cycle slowing will require further exploration.

Finally, how the changing histone landscape contributes to ZGA remains an important open question. We have shown that bulk H3.3 incorporation does not depend on transcription from zelda-dependent genes. However, the reciprocal relationship remains untested. H3.3 incorporation may increase transcription factor accessibility at specific genomic loci to mark them for activation. It is also possible that H3.3 incorporation occurs as a response to transcription initiated by other transcription factors but does not specifically respond to the pioneer factor zelda. We have shown that disruption of major ZGA does not impair bulk H3.3 incorporation, but the role of H3.3 containing nucleosomes in ZGA remains to be tested.

## Materials and methods

### *Drosophila* stocks and genetic crosses

A. y,w; 1xHisC.H3-Dendra2;
B. y,w; H3.3A-Dendra2/CyO;
C. y,w; IX HisC.H3-Dendra2; *shkl^GM163^*/TM3
D. y,w; IX HisC.H3-Dendra2; *shkl^GM130^*/TM3
E. y,w; H3.3A-Dendra2/CyO; *shkl^GM163^*/TM6B
F. y,w; H3.3A-Dendra2/CyO; *shkl^GM130^*/TM6B
G. y,w; H3.3A-Dendra2^S31A^/CyO;
H. y,w; H3.3A-Dendra2^SVM^/CyO;
I. y,w; H3.3A-Dendra2^ASVM^/CyO;
J. *ssm^185b^*,w/FM7c,*w^a^*; IX HisC.H3-Dendra2;
K. *ssm^185b^*,w/FM7c,*w^a^*; H3.3A-Dendra2/CyO;
L. *ssm^185b^*,w/FM7c,*w^a^*; H3.3A-Dendra2^S31A^/CyO;
M. *ssm^185b^*,w/FM7c,*w^a^*; H3.3A-Dendra2^SVM^/CyO;
N. *ssm^185b^*,w/FM7c,*w^a^*; H3.3A-Dendra2^ASVM^/CyO;
O. ) y[1] v[1]; P{y[+t7.7] v[+t1.8]=TRiP.HMJ21114}attP40 (Slbp-RNAi)
P. ;;UAS-Zld-shRNA
Q. yw; Mat-ɑ-tub67-gal4, H3.3A-Dendra2 / CyO; Mat-ɑ-tub15
R. y[1] sc[*] v[1] sev[21]; P{y[+t7.7] v[+t1.8]=TRiP.GL00094}attP2 (white-RNAi)
S. y,w; H3.3A-Dendra2, grp^1^;

Fly stocks A, B, J, and K were generated previously in our lab and are described in Shindo and Amodeo (2019). The Shkl lines y,w; Sp/CyO; *shkl^GM163^*/TM3 and y,w; Sp/CyO; *shkl^GM130^*/TM3 were a generous gift from Stefano Di Talia, Duke University. Stocks C, D, E, and F were generated in the lab by the genetic crossing of these Shkl lines with lines A and B. Stocks L, M, and N were generated by crossing the stocks G, H and I with *ssm^185b^*,w/FM7c,*w^a^*;; (a generous gift from Eric Weischaus). Stocks O and R were obtained from the Bloomington *Drosophila* Stock Center with IDs #51171 and #35573^74^. Stock P was a generous gift from Christine Rushlow, NYU. Stock Q was generated by performing recombination crosses of B with y,w; Mat-ɑ-tub67-gal4; Mat-ɑ-tub15 (a generous gift from Eric Weischaus)^75^. Stock S was generated by performing recombination crosses of B with y,w; grp1/CyO; flies (a generous gift from Eric Weischaus)^70^.

### Drosophila husbandry

All fly stocks were maintained at room temperature, on standard molasses media. The egg lay cages were set up to collect embryos at 25°C (except for the Slbp RNAi flies). Slbp RNAi egg lay cages and associated control w-RNAi cages were set up at 18°C. Embryos from these cages were collected on apple juice agar plates with yeast paste, dechorionated with 50% bleach for up to 2 minutes, and washed twice with dH_2_O. The *ssm^185b^* embryos were collected from *ssm^185b^*/*ssm^185b^* homozygous females. For shkl embryos, the 2 shkl lines were crossed to obtain *shkl*^GM130e^/*shkl*^GM163e^ transheterozygous females and their embryos were imaged. For all the RNAi crosses, males from the gal4 driver line Q, were crossed with virgins from UAS-RNAi lines (O, P or R) to obtain progeny expressing both UAS and Gal4. Embryos from these progeny flies were used for imaging. Embryos from w-RNAi flies were used as controls for all RNAi experiments. Chk1^−/−^ embryos were collected from grp^1^/grp^1^ homozygous females.

### Plasmids and transgenesis

To generate stocks G, H, and I, CRISPR-Cas9 editing was performed at the endogenous H3.3A locus. To this end, pScarlessHD-H3.3A-Dendra2-DsRed plasmid, reported in Shindo and Amodeo (2019)^29^ was modified through site-directed mutagenesis to express H3.3 with H3-specific amino acids, generating pScarlessHD-H3.3A^S31A^-Dendra2-DsRed (S31A mutation), pScarlessHD-H3.3A^SVM^-Dendra2-DsRed (A87S, I89V, G90M mutations) and pScarlessHD-H3.3A^ASVM^-Dendra2-DsRed (S31A, A87S, I89V, G90M mutations) plasmids (Genscript). Two CRISPR target sites were identified using Target Finder^76^, one near the stop codon and one near S31, and the corresponding gRNAs were cloned into pU6-BbsI-chiRNA vector (a gift from Melissa Harrison & Kate O’Connor-Giles & Jill Wildonger, Addgene plasmid #45946). Each mutant plasmid was co-injected with both the gRNA plasmids into nos-Cas9 embryos (TH00787.N) and DsRed+ progeny were selected (BestGene). These progeny were then crossed with nos-PBac flies (a generous gift from Robert Marmion and Stas Shvartsman) to remove the DsRed marker. DsRed negative single males were then crossed with y,w;Sp/CyO; to establish stocks G, H, and I. Insertion of Dendra2 tagged mutants was verified by PCR and Sanger sequencing.

### Microscopy

For live imaging, dechorionated embryos were mounted on glass-bottom MatTek dishes in deionized water and imaged with the 20x, 0.8 NA, objective of Zeiss LSM980 confocal microscope with Airyscan-2 at 45 s intervals for 2 h at room temperature (19-22°C). All H3-Dendra2 tagged embryos were imaged using a 488 nm laser at 2% power and all lines expressing Dendra2 tagged proteins from the endogenous H3.3A locus (H3.3 and the chimeras) were imaged with a 488 nm with 0.5% power in Airyscan multiplex CO-8Y mode. All but shkl embryos and their controls were imaged at a 700 x 700 pixels resolution, with 1 µm Z-steps over a 15 µm range, with a frame time of 26.06ms. All Shkl embryos and their controls (Figure 3, 4, S3) were imaged at a 2836 x 2836 pixels resolution, with 1.2 µm Z-steps over a 14.4 µm range, with a frame time of 328.29ms. All images were acquired with a pixel size of 0.149 µm x 0.149 µm.

### Nuclear export and unbound H3.3 measurement through Dendra2 photoconversion

For measuring the nuclear export and amount of free histone H3.3 (Figure 1E and S1E), we used the photoconvertible Dendra2 tag and the interactive bleaching panel in Zen software. We used a 4µm diameter circular stencil to interactively photo-convert the nuclei. H3.3-Dendra2 within a single nucleus was photoconverted from green to red using a 405 nm laser at 3% power with 60 iterations of laser exposure at a speed of 1.37µs/pixel. The nucleus was converted in the middle of each nuclear cycle for NC11-13 and then imaged with 561 nm at 1% laser power and 488 nm with 0.5% laser power at 15-second intervals until the end of the nuclear cycle. Images were captured at 576 x 576 pixels resolution with 1 µm Z-steps over a 15 µm range, with a frame time of 66.55ms for each channel. The images were acquired with a 40x oil immersion objective, 1.3 NA with a pixel size of 0.092 µm x 0.092 µm.

### Photobleaching corrections

To assess the potential effects of fluorophore photobleaching during our image capture, we performed parallel embryo experiments. In these experiments, we identified 2 embryos of the same age and imaged the interphase nucleus and the metaphase chromatin for both in NC10. Following this, we image only a sub-region of one of the two embryos continuously with our experimental settings described for H3-Dendra2 embryos above until NC13, while keeping the other embryo to develop parallelly without imaging. Once the imaged embryo reached NC13, both the imaged and unimaged parallel embryo were imaged again. We quantified the total nuclear signal from both embryos to evaluate the photobleaching effects. We then compared the continuously imaged section of the embryo, with the area outside the sub-region imaged as well as the unimaged parallel embryo. Using these comparisons, we determined that the effect of photobleaching was minimal and therefore did not apply a numeric photobleaching correction to our data (Figure S1G-H, Table S4-5).

### Nuclear segmentation and intensity analysis

All raw CZI output files from ZEN 3.3 (blue edition) live imaging were first 3D Airyscan Processed at a strength of 3.7 and then converted into individual TIFF files.

For mitotic chromatin quantification, the time points corresponding to metaphase chromatin from each nuclear cycle were extracted and the z-stacks were sum projected in FIJI (2.14.0/1.54f) These files were segmented using the ‘pixel classification + object classification’ applet in the ilastik-1.4.0 software^77^ into chromatin and cytoplasm. The individually segmented mitotic chromatin objects were then exported as a single CSV file containing object properties such as total intensity, mean intensity, and size. The total intensity within each chromatin mass was calculated and normalized to the average NC10 chromatin values (or NC11 for shkl embryos and their controls) for that genotype.

In Shkl embryos and their controls (Figure 3C-F, S3A-B), chromatin was segmented from different regions within an embryo (middle and pole regions for control, and from low and high-density regions for Shkl). In control embryos, middle regions were defined by outlining a box (250×250 pixels) in NC10 around the line separating the embryo into 2 halves. A similar-sized box was outlined with one edge at the tip of the embryo to define the pole region. In shkl embryos, the regions with the highest apparent nuclear density within the center was defined as the high-density region and the region that underwent the partial extra division in NC14 was defined as the low-density region. To account for the asynchronous nature of the divisions in the shkl embryo, within each region, 5-6 nuclei that divided synchronously along the mitotic wave were quantified. For both control and shkl embryos, at least 5 nuclei per embryo were quantified in each cycle for each region.

For analyzing the interphase nuclear concentrations, nuclei from 45 seconds before the nuclear envelope breakdown were segmented in 3D using the ‘pixel classification + object classification’ applet on ilastik software. The CSV file with the mean intensities of each nucleus was exported and normalized to the average NC10 nuclear concentration values for each genotype.

For obtaining the nuclear import curves, individual nuclear cycles were run through the pixel classification + object classification applet in the ilastik software. The results were exported as CSV files and processed with a custom R script. For Shkl embryos (Figure 4B-C,F-G), the pixel prediction maps were used with the ‘tracking with learning’ applet (ilastik) to segment the nuclei as well as track them over time. The tracking result with object properties was exported as a CSV file and processed with a custom R script. Intensities were normalized by the average total intensity of the nuclei at their maximum size in each cycle. For each case, the volume was calculated by multiplying the voxel size with the ‘size in pixels’ of an object.

### Neighborhood analysis

Nuclei within each embryo were tracked over a single nuclear cycle using the ‘tracking + learning’ applet on ilastik. The tracking result with the coordinates of each nucleus over time was obtained as a CSV file, along with other parameters including the total intensity, mean intensity, and nuclear size. The CSV file was analyzed to calculate the number of nuclear neighbors for each nucleus within a 20 µm radius using a custom R script. The script calculates the number of neighbors each nucleus has at its minimum volume since the maximum nuclear import occurs at this time point. To overcome the noise from the incomplete edge nuclei, which are centered lower in the embryo, we utilized the differences in their Z-coordinates to filter them out, after using them for the number of neighbor calculations. For Shkl embryos, as the nuclear cycles are asynchronous, the total intensity traces were aligned to match their minimum volumes to T0. Nuclei with the same number of neighbors were binned together and weighted to reflect the number of nuclei being averaged. The total intensity curves were then normalized such that the average total intensity of the nuclei at their maximum size was equal to 1.

### Cell cycle time measurements

Cell cycle durations were measured from metaphase to metaphase. To account for day-to-day temperature variability, we normalized the mean NC11 durations in control embryos to 10 minutes and scaled for other cell cycles in all embryos acquired on the same day accordingly as done previously^70^.

### Western blot analysis

For western blotting, embryos were collected from white-RNAi flies or Slbp-RNA flies, for a period of 1 hr. Following this the embryos were dechorionated with 50% bleach for 2 minutes followed by 2 washes with deionized water. They were then collected in a microcentrifuge tube and lysed with forceps in ice-cold embryo lysis buffer (50 mM Tris pH 8.0, 150 mM NaCl, 0.5% Triton-X, 1 mM MgCl_2_, 0.1 mM EDTA, 1X protease inhibitor cocktail (Sigma: P2714)). 25 embryos were collected per genotype to quantify pan-H3 levels. Lamelli buffer was added in 1:1 volume and the samples were boiled at 95°C for 5 minutes. The protein lysates were run on a TGX 12% acrylamide gel (Bio-Rad Laboratories), stain-free activated for 45 secs under UV, and transferred onto a LF-PVDF membrane. Membranes were incubated in rabbit anti-H3 antibody (1:1000, Abcam: ab1791) overnight at 4°C. They were then washed and incubated 2 hrs in Alexa Fluor 647-conjugated donkey anti-rabbit IgG antibody (1:2000, Invitrogen: A31573). The membranes were then imaged to detect for fluorescence using a gel imager (Bio-Rad ChemiDoc MP).

### Statistical analysis

Two-way ANOVA tests were conducted to assess the statistical significance between the dataset means of different genotypes over the nuclear cycles. All studies were performed with nuclei from at least 3 embryos. For Shkl embryos, a two-way ANOVA test was used to determine the statistical significance of nuclei within different regions of the same embryo over the different nuclear cycles, with each nucleus as a replicate. For all other embryos, the average chromatin/nuclear values for each NC from each embryo were considered as a replicate. Results from these tests are reported in supplementary tables S1-9.

## Acknowledgements

We thank Shruthi Balachandra, Eric Alpert, Grace Carey and Kiera Schwarz for constructive discussion of this manuscript. We further thank the members of Bickel, Kasper, Lacefield, Moseley and Landino labs at Dartmouth for their helpful suggestions. We thank Patrick Robison from the Dartmouth bioMT Core, Ann Lavanway and Britton Johnson for technical support. We thank Stefano Di Talia for shkl stock flies, Eric Weischaus for ssm, grp and gal4-driver stock flies; Robert Marmion for the nos-PBac transposase flies, Christine Rushlow for the zelda-RNAi flies and Robert Duronio for the 1xHisC plasmid. We thank FlyBase, funded by NHGRI and NIGMS, and Bloomington Drosophila Stock Center (NIHP40OD018537) for providing essential resources. This work was funded by NIH/NIGMS (P20-GM113132 and R35GM150853 to AAA). ADB is supported by Sondra and Charles Gilman Graduate Research Fellowship.

## Author contributions

Conceptualization, ADB, MGB, YS and AAA.; Investigation and Analysis, ADB, MGB and ABW; Writing – Original Draft, ADB, MGB and AAA; Writing – Review & Editing, ADB, MGB, YS and AAA.

