## Supplemental materials for "Local nuclear to cytoplasmic ratio regulates H3.3 incorporation via cell cycle state during zygotic genome activation"

**Supplementary Figure 1:Tools and controls underlying image quantification of mitotic and interphase Dendra2 and H3.3 export**

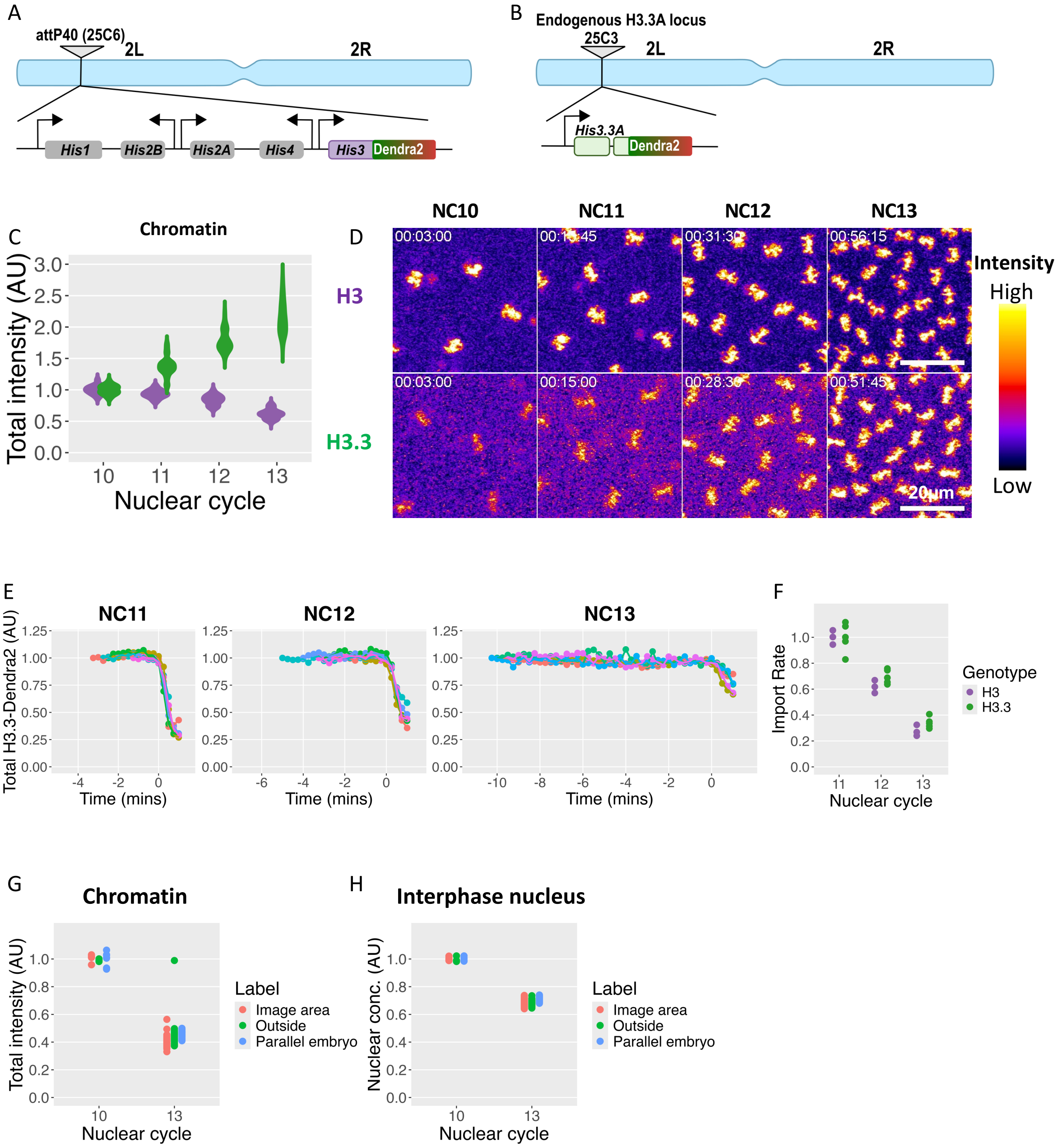


### Supplementary Figure 1: Tools and controls underlying image quantification of mitotic and interphase Dendra2 and H3.3 export

**(A)** A 5kb region containing a single histone gene cluster with one copy of each of the 5 replication-coupled histones including, their promoters, and UTRs as in the endogenous locus in which H3 was N-terminally tagged with the green-to-red photo-switchable fluorophore Dendra2 was inserted at the attp40 site on chromosome 2L (from Shindo and

Amodeo, 2019). **(B)** The endogenous H3.3A gene locus was edited to express an N-terminally tagged H3.3-Dendra2 using CRISPR/Cas9 (from Shindo and Amodeo, 2019). **(C)** Total pixel intensities (corresponding to total amounts) on mitotic chromatin for H3-Dendra2 (purple) and H3.3-Dendra2 (green) between NC10-13 normalized to the average individual NC10 values. Chromatin-bound H3 decreases over NC10-13 whereas H3.3 increases. **(D)** Maximum intensity projections of H3-Dendra2 (top) and H3.3-Dendra2 (bottom) on mitotic chromatin from NC10-13. Images are pseudo-colored with non-linear look-up tables such that purple indicates low intensities and yellow indicates high intensities. H3 intensities fall and H3.3 intensities rise over the cycles. **(E)** Initial slopes of the nuclear import curves shown in 1C and 1D for NC11-13. All slopes are normalized to NC11 values. **(F)** The total pixel intensity of individual photoconverted H3.3-Dendra2 nuclei in NC11-13 (n = 5). Time is shown relative to nuclear envelope breakdown (NEB). H3.3-Dendra2 intensity remains constant before NEB indicating that the nuclear export is negligible. The loss of red signal at NEB represents the pool of unbound H3.3 in the nucleus. Each trace represents a single nucleus, each from different embryos. These data were used to plot the unbound H3.3 fraction in Figure 1E. **(G-H)** Total pixel intensity of H3-Dendra2 on chromatin **(G)** and interphase nuclei **(H)** with and without continued laser exposure. Two parallel embryos were used to obtain NC10 and NC13 mitotic chromatin with different levels of laser exposure for photobleaching correction (see methods for details). The data were divided into 3 regions: ‘Image area’ corresponds to nuclei imaged throughout NC10-13; ‘outside’ corresponds to nuclei outside the image area, but within the imaged embryo; and ‘parallel embryo’ corresponds to nuclei that were only imaged once in NC10 and once in NC13 without continued exposure. Photobleaching was observed to be negligible in experiments quantifying both the mitotic chromatin and interphase nuclear concentration. (Statistical comparisons for C, E, G and H can be found in Supplemental tables 1,3-5).

**Supplementary Figure 2: Representative images of H3/H3.3 chimeras, import curves**

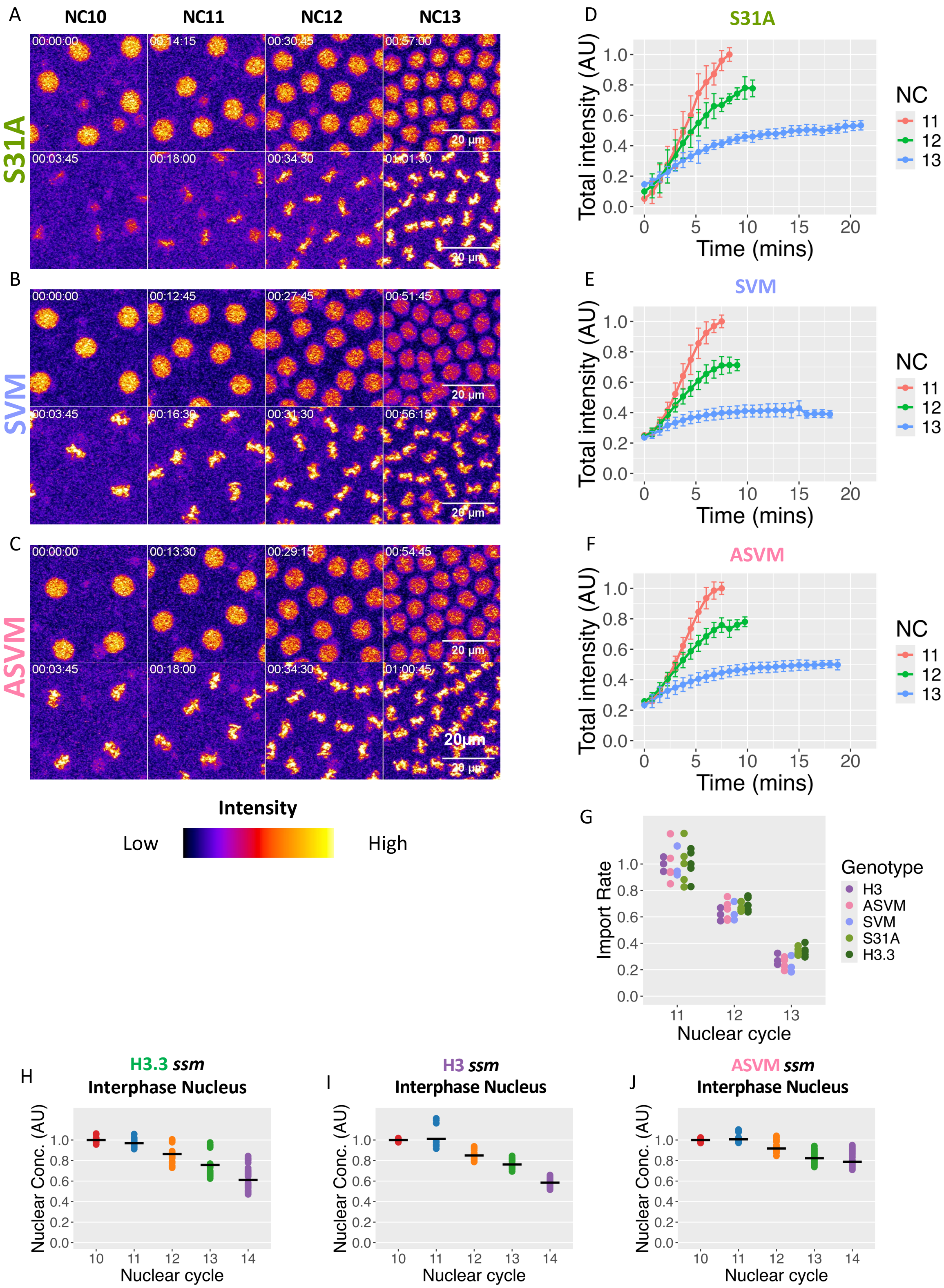

**Supplementary Figure 2: Representative images of H3/H3.3 chimeras, import curves**

**(A)** Representative maximum intensity projections of H3.3$^{S31A}$-Dendra2 during interphase and mitosis over NC10-13: interphase nuclei (top) and mitotic chromatin (bottom). Images are pseudo-colored with non-linear look-up tables such that purple indicates low intensities and yellow indicates high intensities. H3.3$^{S31A}$ behaves similarly to H3.3. **(B)** Representative maximum intensity projections of H3.3$^{SVM}$-Dendra2 during interphase and mitosis over NC10-13: interphase nuclei (top) and mitotic chromatin (bottom). H3.3$^{SVM}$ behaves similarly to H3. **(C)** Representative maximum intensity projections of H3.3$^{ASVM}$-Dendra2 during interphase and mitosis over NC10-13: interphase nuclei (top) and mitotic chromatin (bottom). H3.3$^{ASVM}$ behaves similarly to H3. Data from embryos in A-C are quantified in Figure 2. **(D-F)** Total pixel intensities over time for NC11-13 normalized to the maximum NC11 values for H3.3$^{S31A}$-Dendra2 **(D)**, H3.3$^{SVM}$-Dendra2 **(E)**, and H3.3$^{ASVM}$-Dendra2 **(F)**. H3.3$^{S31A}$-Dendra2 import is similar to H3.3-Dendra2, and only slows after 5 minutes without plateauing. H3.3$^{SVM}$-Dendra2 and H3.3$^{ASVM}$-Dendra2 import in a similar manner to H3-Dendra2 and plateau after 5 minutes. **(G)** The initial slopes of nuclear import curves of chimeras shown in D-F for NC11-13. H3-Dendra2 and H3.3-Dendra2 slopes from S1E are included for reference. All slopes are normalized to NC11 values. **(H-J)** Average interphase nuclear intensities of H3.3-Dendra2 **(H)**, H3-Dendra2 **(I)** and H3.3$^{ASVM}$-Dendra2 **(J)** in Hira$^{ssm}$ embryos 45 seconds before the NEB in NC10-14, normalized to their average intensities in NC10. Though H3.3 is not incorporated, it is imported into the nucleus and its concentrations reduce with each cycle. H3 nuclear concentrations also drop with each cycle. However, H3.3$^{ASVM}$ concentrations are relatively more stable over the cycles. (n=5 all chimeras, 3 H3 ssm, 4 H3.3 ssm, and 5 H3.3$^{ASVM}$ ssm embryos. Statistical comparisons for G can be found in Supplemental table 8).

**Supplementary Figure 3: Replicate shkl embryos demonstrate consistent effects**

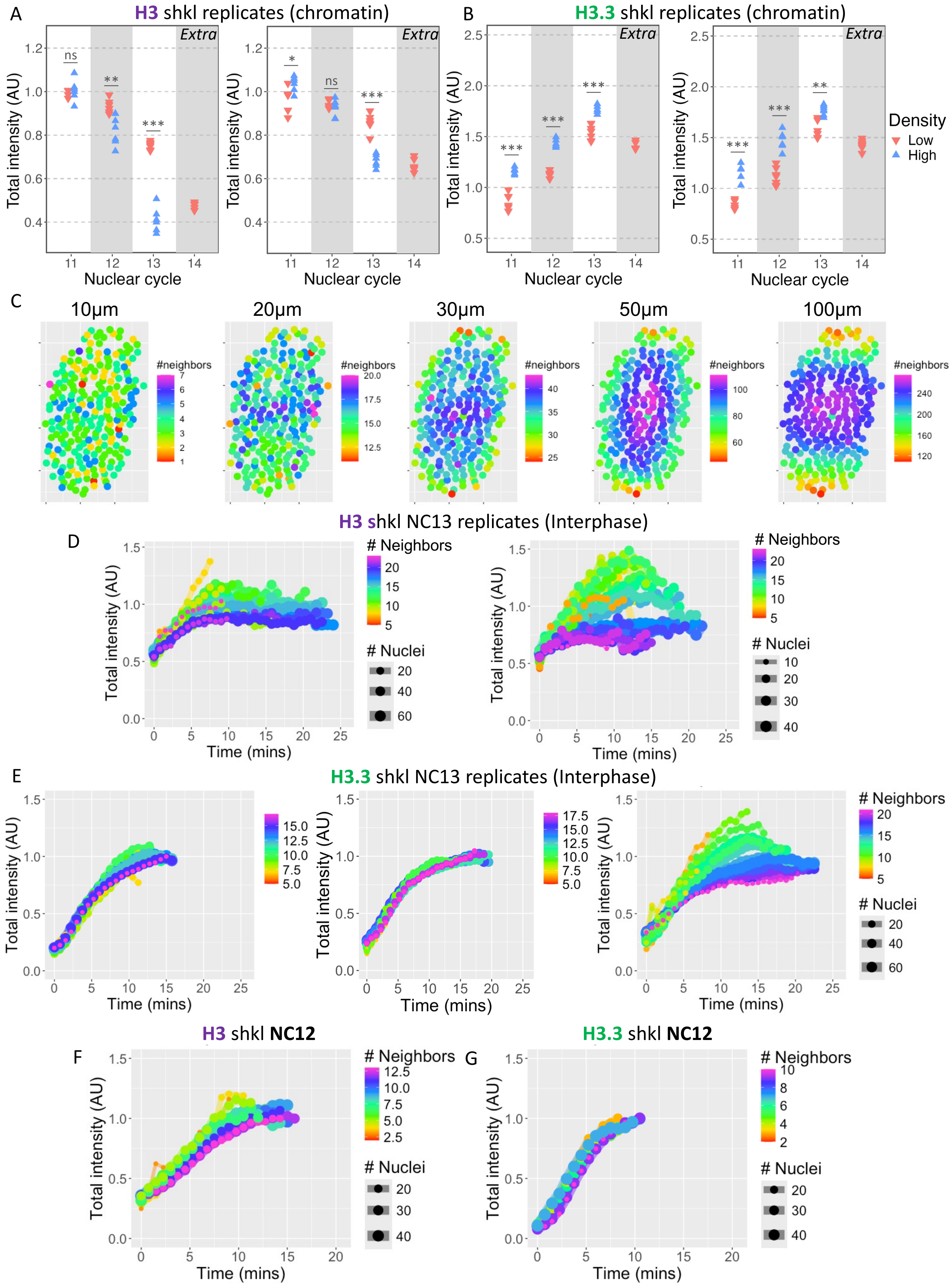

**Supplementary Figure 3: Replicate shkl embryos demonstrate consistent effects**

**(A)** Replicate shkl embryos of the same genotype as shown in Figure 3E demonstrate that H3-Dendra2 is consistently retained on mitotic chromatin by nuclei at low-density regions compared to high-density regions within the same cell cycle over NC11-14. **(B)** Replicate shkl embryos of the same genotype as shown in Figure 3F demonstrate that H3.3-Dendra2 incorporation is reduced on mitotic chromatin by nuclei at low-density regions compared to high-density regions within the same cell cycle over NC11-14. **(C)** Example control embryo in which the radius used to determine the number of neighbors was varied from 10 to 100 μm as shown. A 20 μm radius was deemed optimal for neighborhood analysis as it enabled us to accurately capture the gradient observed in the shkl embryos while excluding edge effects due to embryo curvature. **(D)** Replicate shkl embryos of the same genotype as shown in Figure 4F demonstrate that nuclear import and accumulation of H3 inversely correlate with the number of neighbors surrounding a given nucleus, suggesting H3 nuclear import is N/C ratio sensitive. **(E)** Replicate shkl embryos of the same genotype as shown in Figure 4G demonstrate that nuclear import and accumulation of H3.3 is less N/C ratio sensitive than H3 in most cases. **(F-G)** Total intensities over time for H3-Dendra2 **(F)** and H3.3-Dendra2 **(G)** in NC12 shkl embryos indicate that the trends observed in NC13 begin in NC12 though to a lesser extent. These data were taken from the same embryo that was used in Figure 4F and G respectively. (Statistical significance was determined by 2-way ANOVA, ns= $p>.05$, *=$p<.05$, **=$p<0.01$, *** = $p<0.001$)

**Supplementary Figure 4: Cell cycle times in mutant embryos, H3 levels in SLBP RNAi lines**

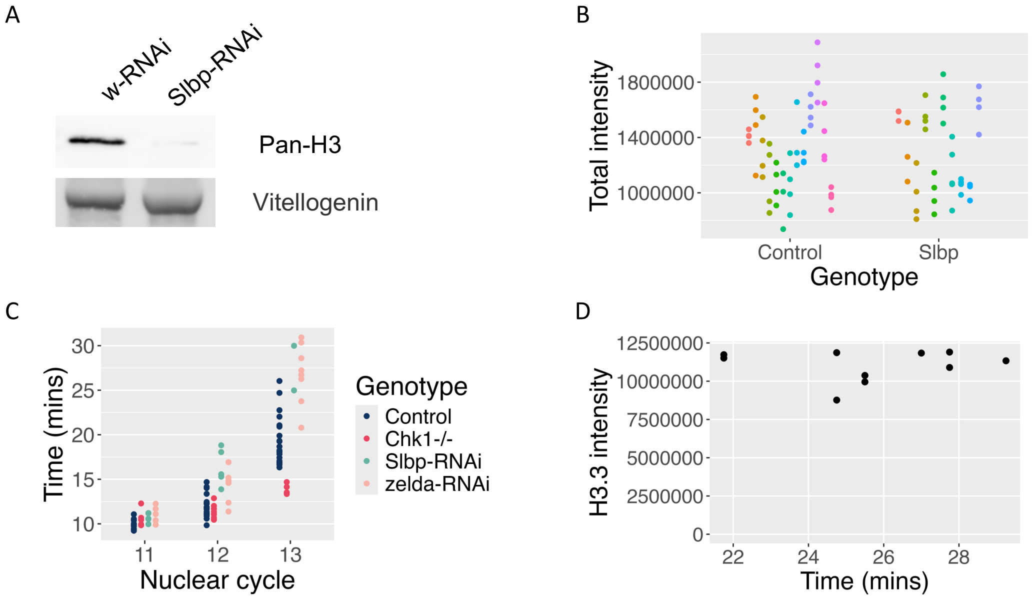


**Supplementary Figure 4: Cell cycle times in mutant embryos, H3 levels in SLBP RNAi lines**

**(A)** Western blot against a Pan-H3 antibody in w-RNAi (control) and Slbp-RNAi embryos after a 1hr collection. Stainfree signal for Vitellogenin (~45kDa) is used as a loading control. Slbp-RNAi causes H3 knockdown. **(B)** H3.3 incorporation in NC10 mitotic chromatin in w-RNAi (control) and severely affected Slbp embryos, which do not survive past NC11 due to mitotic defects. H3.3 levels are comparable between w-RNAi (control) and Slbp embryos. Different colors represent nuclei from different embryos. **(C)** Cell cycle durations of control (a mix of y,w;; and w-RNAi) and RNAi/mutant embryos used in Figure 5. Note that cell cycles are shortened in Chk1$^{-/-}$ embryos and lengthened in Slbp-RNAi and zelda-RNAi embryos. **(D)** H3.3 chromatin incorporation versus NC13 duration in control embryos. No correlation is observed between NC13 cell cycle duration and H3.3 incorporation. (n≥5 embryos. Statistical comparisons for C can be found in Supplemental table 9)

**Supplementary movies**

**Movie S1:** Control embryos have uniformly distributed nuclei that divide synchronously. (nuclear marker H3-Denrdra2, Scalebar 50µm, same embryo as Figure 3A).

**Movie S2:** shkl embryos have non-uniformly distributed nuclei with low nuclear density at the posterior pole. Nuclei divide asynchronously. (nuclear marker H3.3-Denrdra2, Scalebar 50µm, same embryo as Figure 3B)

### Supplementary tables

**Supplementary Table S1:** 2-way ANOVA results of H3-Dendra2 and H3.3-Dendra2 on chromatin over NC10 - NC13 as shown in Figure S1C.

| Comparison groups | Difference in means | Adjusted p-value |
|---|---:|---:|
| NC11_H3 - NC10_H3 | -0.0627471 | 0.98843294 |
| NC12_H3 - NC11_H3 | -0.097712 | 0.88298162 |
| NC13_H3 - NC12_H3 | -0.220603 | 0.08782752 |
| NC11_H3.3 - NC10_H3.3 | 0.37315126 | 0.00040692 |
| NC12_H3.3 - NC11_H3.3 | 0.39265816 | 0.00019202 |
| NC13_H3.3 - NC12_H3.3 | 0.37344713 | 0.00040232 |

**Supplementary Table S2:** 2-way ANOVA results of H3-Dendra2 and H3.3-Dendra2 nuclear concentrations in the interphase nucleus during NC10 - NC13 as shown in Figure 1B.

| Comparison groups | Difference in means | Adjusted p-value |
|---|---:|---:|
| NC11_H3 - NC10_H3 | -0.1408876 | 0.18475814 |
| NC12_H3 - NC11_H3 | -0.1115503 | 0.44067226 |
| NC13_H3 - NC12_H3 | -0.1364362 | 0.21426967 |
| NC11_H3.3 - NC10_H3.3 | -0.0241768 | 0.99871742 |
| NC12_H3.3 - NC11_H3.3 | -0.077933 | 0.56567533 |
| NC13_H3.3 - NC12_H3.3 | -0.0730441 | 0.63966091 |

**Supplementary Table S3:** 2-way ANOVA results of initial H3-Dendra2 and H3.3-Dendra2 import rates during NC11 - NC13 as shown in Figure S1E.

| Comparison groups | Difference in means | Adjusted p-value |
|---|---:|---:|
| NC12_H3 - NC11_H3 | -0.381055416 | 0.00016788 |
| NC12_H3.3 - NC11_H3.3 | -0.303255393 | 0.00010136 |
| NC13_H3 - NC12_H3 | -0.34119875 | 0.00110214 |
| NC13_H3.3 - NC12_H3.3 | -0.356357728 | 3.46E-06 |
| NC12_H3.3 - NC12_H3 | 0.077800023 | 0.99461495 |
| NC13_H3.3 - NC13_H3 | 0.062641045 | 0.99943292 |

**Supplementary Table S4:** 2-way ANOVA results for the different regions in the parallel embryo experiment for photobleaching correction for chromatin (NC10 vs NC13) as shown in Figure S1G.

| Comparison groups | Difference in means | Adjusted p-value |
|---|---|---|
| NC10_Outside - NC10_Image area | -0.015039 | 0.99820702 |
| NC10_Parallel embryo - NC10_Image area | -0.006684 | 0.99993379 |
| NC10_Parallel embryo - NC10_Outside | 0.00835498 | 0.99985763 |
| NC13_Outside - NC13_Image area | 0.05703447 | 2.35E-06 |
| NC13_Parallel embryo - NC13_Image area | 0.05682858 | 2.58E-06 |
| NC13_Parallel embryo - NC13_Outside | -0.0002059 | 1 |
| NC13_Image area - NC10_Image area | -0.6050886 | 0 |
| NC13_Outside - NC10_Outside | -0.5330152 | 0 |
| NC13_Parallel embryo - NC10_Parallel embryo | -0.5415761 | 0 |

**Supplementary Table S5:** 2-way ANOVA results for the different regions in the parallel embryo experiment for photobleaching correction for interphase nuclear concentration (NC10 vs NC13) as shown in Figure S1H.

| Comparison groups | Difference in means | Adjusted p-value |
|---|---|---|
| NC10_Outside - NC10_Image area | -0.0030836 | 0.99990859 |
| NC10_Parallel embryo - NC10_Image area | -0.0011564 | 0.99999823 |
| NC10_Parallel embryo - NC10_Outside | 0.00192728 | 0.99998957 |
| NC13_Outside - NC13_Image area | -0.004531 | 0.6781679 |
| NC13_Parallel embryo - NC13_Image area | 0.02075448 | 1.31E-11 |
| NC13_Parallel embryo - NC13_Outside | 0.02528553 | 1.73E-12 |
| NC13_Image area - NC10_Image area | -0.3123329 | 9.00E-14 |
| NC13_Outside - NC10_Outside | -0.3137803 | 9.00E-14 |
| NC13_Parallel embryo - NC10_Parallel embryo | -0.2904221 | 9.00E-14 |

**Supplementary Table S6:** 2-way ANOVA results of chimeras compared to H3-Dendra2 and H3.3-Dendra2 on chromatin over NC10 - NC13 as shown in Figure 2B.

| Comparison groups | Difference in means | Adjusted p-value |
|---|---|---|
| NC10_H3 - NC10_ASVM | -1.55E-15 | 1 |
| NC10_H3.3 - NC10_ASVM | 3.33E-16 | 1 |
| NC10_S31A - NC10_ASVM | 0 | 1 |

| | | |
|---|---|---|
| NC10_SVM - NC10_ASVM | -1.11E-16 | 1 |
| NC10_H3.3 - NC10_H3 | 1.89E-15 | 1 |
| NC10_S31A - NC10_H3 | 1.55E-15 | 1 |
| NC10_SVM - NC10_H3 | 1.44E-15 | 1 |
| NC10_S31A - NC10_H3.3 | -3.33E-16 | 1 |
| NC10_SVM - NC10_H3.3 | -4.44E-16 | 1 |
| NC10_SVM - NC10_S31A | -1.11E-16 | 1 |
| NC11_H3 - NC11_ASVM | -0.002548004 | 1 |
| NC11_H3.3 - NC11_ASVM | 0.43335039 | 2.05E-06 |
| NC11_S31A - NC11_ASVM | 0.195753042 | 0.32719825 |
| NC11_SVM - NC11_ASVM | -0.055004006 | 0.99999902 |
| NC11_H3.3 - NC11_H3 | 0.435898394 | 1.74E-06 |
| NC11_S31A - NC11_H3 | 0.198301045 | 0.30496131 |
| NC11_SVM - NC11_H3 | -0.052456002 | 0.99999955 |
| NC11_S31A - NC11_H3.3 | -0.237597348 | 0.08064289 |
| NC11_SVM - NC11_H3.3 | -0.488354396 | 6.01E-08 |
| NC11_SVM - NC11_S31A | -0.250757047 | 0.04722605 |
| NC12_H3 - NC12_ASVM | 0.007934945 | 1 |
| NC12_H3.3 - NC12_ASVM | 0.934203465 | 0 |
| NC12_S31A - NC12_ASVM | 0.527625335 | 4.57E-09 |
| NC12_SVM - NC12_ASVM | -0.11622197 | 0.97313608 |
| NC12_H3.3 - NC12_H3 | 0.92626852 | 0 |
| NC12_S31A - NC12_H3 | 0.51969039 | 7.72E-09 |
| NC12_SVM - NC12_H3 | -0.124156915 | 0.95000571 |
| NC12_S31A - NC12_H3.3 | -0.40657813 | 1.08E-05 |
| NC12_SVM - NC12_H3.3 | -1.050425436 | 0 |
| NC12_SVM - NC12_S31A | -0.643847306 | 0 |
| NC13_H3 - NC13_ASVM | -0.038619561 | 1 |
| NC13_H3.3 - NC13_ASVM | 1.481699114 | 0 |
| NC13_S31A - NC13_ASVM | 0.861241303 | 0 |
| NC13_SVM - NC13_ASVM | -0.139872287 | 0.8684167 |
| NC13_H3.3 - NC13_H3 | 1.520318675 | 0 |
| NC13_S31A - NC13_H3 | 0.899860864 | 0 |
| NC13_SVM - NC13_H3 | -0.101252726 | 0.99386895 |
| NC13_S31A - NC13_H3.3 | -0.620457811 | 0 |
| NC13_SVM - NC13_H3.3 | -1.621571401 | 0 |
| NC13_SVM - NC13_S31A | -1.00111359 | 0 |

**Supplementary Table S7:** 2-way ANOVA results for nuclear concentrations of chimeras compared to H3-Dendra2 and H3.3-Dendra2 in the interphase nucleus during NC10 - NC13 as shown in Figure 2C.

| Comparison groups | Difference in Means | Adjusted p-value |
|---|---|---|
| NC10_H3 - NC10_ASVM | -0.0055188 | 1 |
| NC10_H3.3 - NC10_ASVM | -0.0055188 | 1 |
| NC10_S31A - NC10_ASVM | 0.00028757 | 1 |
| NC10_SVM - NC10_ASVM | -0.0055188 | 1 |
| NC10_H3.3 - NC10_H3 | 7.94E-11 | 1 |
| NC10_S31A - NC10_H3 | 0.00580633 | 1 |
| NC10_SVM - NC10_H3 | -1.06E-11 | 1 |
| NC10_S31A - NC10_H3.3 | 0.00580633 | 1 |
| NC10_SVM - NC10_H3.3 | -9.00E-11 | 1 |
| NC10_SVM - NC10_S31A | -0.0058063 | 1 |
| NC11_H3 - NC11_ASVM | 0.00619749 | 1 |
| NC11_H3.3 - NC11_ASVM | 0.12290828 | 0.17906869 |
| NC11_S31A - NC11_ASVM | 0.04818638 | 0.99931011 |
| NC11_SVM - NC11_ASVM | 0.00128513 | 1 |
| NC11_H3.3 - NC11_H3 | 0.11671079 | 0.50685044 |
| NC11_S31A - NC11_H3 | 0.0419889 | 0.99998836 |
| NC11_SVM - NC11_H3 | -0.0049124 | 1 |
| NC11_S31A - NC11_H3.3 | -0.0747219 | 0.91786474 |
| NC11_SVM - NC11_H3.3 | -0.1216232 | 0.19276663 |
| NC11_SVM - NC11_S31A | -0.0469013 | 0.99952054 |
| NC12_H3 - NC12_ASVM | 0.0202021 | 1 |
| NC12_H3.3 - NC12_ASVM | 0.1705302 | 0.00518502 |
| NC12_S31A - NC12_ASVM | 0.0328656 | 0.99999765 |
| NC12_SVM - NC12_ASVM | -0.0434382 | 0.99983517 |
| NC12_H3.3 - NC12_H3 | 0.1503281 | 0.1149165 |
| NC12_S31A - NC12_H3 | 0.0126635 | 1 |
| NC12_SVM - NC12_H3 | -0.0636403 | 0.99621661 |
| NC12_S31A - NC12_H3.3 | -0.1376646 | 0.06981896 |
| NC12_SVM - NC12_H3.3 | -0.2139684 | 8.38E-05 |
| NC12_SVM - NC12_S31A | -0.0763038 | 0.90297737 |
| NC13_H3 - NC13_ASVM | 0.00965776 | 1 |
| NC13_H3.3 - NC13_ASVM | 0.22337804 | 3.21E-05 |
| NC13_S31A - NC13_ASVM | 0.06052475 | 0.9889634 |
| NC13_SVM - NC13_ASVM | -0.0863012 | 0.77259947 |
| NC13_H3.3 - NC13_H3 | 0.21372028 | 0.00139254 |

| | | |
|---|---|---|
| NC13_S31A - NC13_H3 | 0.05086699 | 0.99979886 |
| NC13_SVM - NC13_H3 | -0.095959 | 0.82075511 |
| NC13_S31A - NC13_H3.3 | -0.1628533 | 0.01001226 |
| NC13_SVM - NC13_H3.3 | -0.3096793 | 2.76E-09 |
| NC13_SVM - NC13_S31A | -0.146826 | 0.03597147 |

**Supplementary Table S8:** 2-way ANOVA results of initial import rates of chimeras compared to H3-Dendra2 and H3.3-Dendra2 during NC11 - NC13 as shown in Figure S2G.

| Comparison groups | Difference in Means | Adjusted p-value |
|---|---|---|
| NC12_ASVM - NC11_ASVM | -0.346793281 | 6.39E-06 |
| NC12_H3 - NC11_H3 | -0.381055416 | 0.00016788 |
| NC12_H3.3 - NC11_H3.3 | -0.303255393 | 0.00010136 |
| NC12_S31A - NC11_S31A | -0.320997327 | 3.31E-05 |
| NC12_SVM - NC11_SVM | -0.363066437 | 0.0003963 |
| NC13_ASVM - NC12_ASVM | -0.398034537 | 2.36E-07 |
| NC13_H3 - NC12_H3 | -0.34119875 | 0.00110214 |
| NC13_H3.3 - NC12_H3.3 | -0.356357728 | 3.46E-06 |
| NC13_S31A - NC12_S31A | -0.341763381 | 8.82E-06 |
| NC13_SVM - NC12_SVM | -0.400466562 | 6.55E-05 |
| NC13_SVM - NC13_ASVM | -0.018705181 | 1 |
| NC13_SVM - NC13_H3 | -0.041278832 | 0.99999904 |
| NC12_ASVM - NC12_H3 | 0.034262135 | 0.99999964 |
| NC12_H3.3 - NC12_ASVM | 0.043537888 | 0.99995511 |
| NC12_H3.3 - NC12_H3 | 0.077800023 | 0.99461495 |
| NC12_H3.3 - NC12_S31A | 0.017741934 | 1 |
| NC12_H3.3 - NC12_SVM | 0.059811044 | 0.99966109 |
| NC12_S31A - NC12_ASVM | 0.025795954 | 0.99999994 |
| NC12_S31A - NC12_H3 | 0.060058089 | 0.99964501 |
| NC12_S31A - NC12_SVM | 0.04206911 | 0.99999497 |
| NC12_SVM - NC12_ASVM | -0.016273156 | 1 |
| NC12_SVM - NC12_H3 | 0.017988979 | 1 |
| NC13_ASVM - NC13_H3 | -0.022573651 | 1 |
| NC13_H3.3 - NC13_ASVM | 0.085214697 | 0.95767702 |
| NC13_H3.3 - NC13_H3 | 0.062641045 | 0.99943292 |
| NC13_H3.3 - NC13_S31A | 0.003147587 | 1 |
| NC13_H3.3 - NC13_SVM | 0.103919878 | 0.93613851 |
| NC13_S31A - NC13_ASVM | 0.082067109 | 0.96862842 |

| | | |
|---|---|---|
| NC13_S31A - NC13_H3 | 0.059493458 | 0.99968081 |
| NC13_S31A - NC13_SVM | 0.100772291 | 0.94917367 |

**Supplementary Table S9:** 2-way ANOVA results of cell cycle durations for RNAi/mutant embryos as shown in Figure S4B.

| Comparison groups | Difference in means | Adjusted p-value |
|---|---|---|
| NC11_Slbp-RNAi - NC11_Control | 0.69586667 | 0.99950082 |
| NC12_Slbp-RNAi - NC12_Control | 4.4048 | 3.30E-05 |
| NC13_Slbp-RNAi - NC13_Control | 8.49575 | 2.35E-08 |
| NC11_zelda-RNAi - NC11_Control | 0.87616667 | 0.97411098 |
| NC12_zelda-RNAi - NC12_Control | 2.45433333 | 0.01554046 |
| NC13_zelda-RNAi - NC13_Control | 8.03791667 | 7.69E-14 |
| NC11_Chk1$^{-/-}$ - NC11_Control | 0.42016667 | 0.99997836 |
| NC12_Chk1$^{-/-}$ - NC12_Control | -0.5474091 | 0.99912207 |
| NC13_Chk1$^{-/-}$ - NC13_Control | -5.05825 | 1.13E-05 |
